# MeCP2 gates spatial learning-induced alternative splicing events in the mouse hippocampus

**DOI:** 10.1101/2020.06.25.170480

**Authors:** David V.C. Brito, Kubra Gulmez Karaca, Ana M.M. Oliveira

## Abstract

Long-term memory formation is supported by functional and structural changes of neuronal networks, which rely on *de novo* gene transcription and protein synthesis. The modulation of the neuronal transcriptome in response to learning depends on transcriptional and post-transcriptional mechanisms. DNA methylation writers and readers regulate the activity-dependent genomic program required for memory consolidation. The most abundant DNA methylation reader, the Methyl CpG binding domain protein 2 (MeCP2), has been shown to regulate alternative splicing, but whether it establishes splicing events important for memory consolidation has not been investigated. In this study, we identified the alternative splicing profile of the mouse hippocampus in basal conditions and after a spatial learning experience, and investigated the requirement of MeCP2 for these processes. We observed that spatial learning triggers a wide-range of alternative splicing events in transcripts associated with structural remodeling and that virus-mediated knockdown of MeCP2 impairs learning-dependent post-transcriptional responses of mature hippocampal neurons. Furthermore, we found that MeCP2 preferentially affected the splicing modalities intron retention and exon skipping and guided the alternative splicing of distinct set of genes in baseline conditions and after learning. Lastly, comparative analysis of the MeCP2-regulated transcriptome with the alternatively spliced mRNA pool, revealed that MeCP2 disruption alters the relative abundance of alternatively spliced isoforms without affecting the overall mRNA levels. Overall our findings reveal that adult hippocampal MeCP2 is required to finetune alternative splicing events in basal conditions, as well as in response to spatial learning. This study provides new insight into how MeCP2 regulates brain function, particularly cognitive abilities, and sheds light onto the pathophysiological mechanisms of Rett syndrome, that is characterized by intellectual disability and caused by mutations in the *Mecp2* gene.

## Introduction

It is well established that long-term memory formation requires *de novo* gene transcription and protein synthesis. In response to neuronal activity, immediate early genes are rapidly transcribed, many of which initiate a second, delayed wave of gene transcription [1]. These newly synthesized mRNAs and proteins guide neuronal structural and functional changes that support memory consolidation [1, 2]. This complex process is regulated at multiple layers by the action of transcription factors and epigenetic players as well as chromatin architecture organizers that alter the accessibility of transcribed loci [3–7]. At the same time, neuronal activity triggers alternative splicing events that offer another level of gene expression regulation. Indeed, several studies have reported activity-dependent alternative splicing mechanisms in neurons [8–12] whose disruptions are associated with brain disorders [13]. Furthermore, selective expression of alternative splice variants functionally impacts the cells through the remodeling of the transcriptome which may modify protein interaction, function and localization [10, 14–16]. Altogether, these findings strongly suggest that the coordinated regulation of gene transcription and alternative splicing is vital to determine neuronal activity-dependent changes required for memory consolidation.

DNA methylation is a dynamically regulated epigenetic mark that controls activity-dependent transcription and alternative splicing [5, 17]. Methyl CpG binding domain protein 2 (MeCP2) is the most abundant DNA methylation reader in the brain, linking DNA methyl marks to higher order chromatin architectural changes through interaction with its numerous binding partners [18, 19]. MeCP2 function is essential during neurodevelopment, since reduction in MeCP2 levels culminates in a severe neurological disorder, Rett Syndrome (RTT) [20]. Similarly, MeCP2 is indispensable during adulthood; it gates adult cognitive abilities and maintains chromatin architecture and proper functioning of the brain [18, 21]. Until now, MeCP2 has been repeatedly shown to impact the transcriptional profile of developing and mature neurons in basal conditions, as well as in response to neuronal activity [22, 23]. In contrast, less is known about its functions in alternative splicing mechanisms regulating synaptic plasticity changes required for the formation and maintenance of memory. Recent studies identified that MeCP2 interacts with alternative splicing components, (e.g., Y-box binding protein 1 [YB-1]), and regulates their expression to influence alternative splicing events in neuroblastoma [24] or cancer cell lines [25]. Reduced DNA methylation that leads to reduced binding of MeCP2 to DNA was shown to decrease alternative splicing events and increased intron-retention mechanisms in embryonic stem cells, and in human and mouse cell lines [26]. Moreover, in mouse models of RTT, MeCP2 was shown to control alternative splicing events in the cortex during basal conditions [24, 27] and in the hippocampus in basal state and in a seizure model [8]. Altogether these studies have attributed a role for MeCP2 in the regulation of alternative splicing, however it remains unclear whether MeCP2 establishes alternative splicing events important for memory consolidation in response to a physiological learning stimulus.

Therefore, in this study we aimed to investigate the alternative splicing regulatory function of MeCP2 in the adult hippocampus of mice during spatial memory consolidation. We used a previously characterized mouse model in which MeCP2 is selectively reduced in the adult hippocampus to exclude possible confounds originating from impaired development or from abnormal anxiety- and motor abilities [21]. Using this model, we performed RNA-seq to determine genome-wide alternative splicing events regulated by MeCP2 in basal conditions, as well as after a non-aversive spatial learning task. We identified a novel set of learning-induced alternative splicing variants in the mouse hippocampus. Furthermore, we found that MeCP2 knockdown altered the alternatively spliced mRNA profile of hippocampal neurons in basal conditions and abolished the alternative splicing events triggered by learning, mostly affecting exon-exclusion and intron-retention mechanisms involved in synaptic plasticity. Moreover, by comparative analysis of MeCP2-regulated transcriptome with alternatively spliced mRNA pool, we provided evidence that MeCP2 knockdown altered the relative abundance of alternatively spliced mRNAs even if the overall levels of the gene were not changed. Overall, our results attribute a novel role to MeCP2 in guiding basal and learning-induced alternative splicing mechanisms in mature hippocampal neurons required for long-term memory formation.

## Materials and Methods

### Mice

Throughout the study, we used adult male C57BL/6N mice that were 8 weeks old at the time of surgery [(MeCP2-shRNA (n=8) or Control-shRNA (n=8)] (Charles River, Sulzfeld, Germany). The mice were group-housed on a 12h light/dark cycle with *ad libitum* access to food and water. All behavioral experiments were carried during the light phase. Mice that were sick and/or injured from cage-mate fighting were not included into the study. All procedures were performed according to the German guidelines for the care and use of laboratory animals and with the European Community Council Directive 86/609/EEC.

### Recombinant adeno-associated virus (rAAV)

Viral particles were produced and purified as described previously [28]. For the knockdown of MeCP2, we used a vector containing the U6 promoter upstream of the short-hairpin RNA (shRNAs) (MeCP2-specific or control) sequence and a CamKIIα promoter driving mCherry expression (as an infection control) as described previously [21].

### Stereotaxic surgery

A total volume of 1.5 μl (2:1 mixture of rAAV solution and 20% mannitol) was injected into the dorsal hippocampus (relative to Bregma: −2 mm anteroposterior, ±1.5 mm mediolateral, −1.7, −1.9 and −2.1 mm dorsoventral) at the speed of 200 nl/min as previously described [21, 29].

### Spatial-object recognition task

Spatial-object recognition task was executed as explained previously [30]. Briefly, after mice were habituated to the experimenter and behavioral room by gentle handling (3 consecutive days, 1 min per mouse), mice were placed into an open arena (50 cm×50 cm×50 cm with a visual cue placed on the arena wall) in the absence of objects. In the next three sessions, mice were able to explore two distinct objects (a glass bottle and a metal tower). Between the sessions, mice were placed in their home cage for 3 min.

### RNA-Sequencing

30 min after training in spatial object recognition task, the infected dorsal hippocampal tissue (identified by mCherry expression) was micro-dissected for RNA-seq analysis. Home-cage mice were not subjected to training, but dissected simultaneously with trained mice to account for time of the day differences. Total RNA was isolated using the RNeasy Plus Mini Kit (Qiagen, Hilden, Germany) with additional on-column DNase I digestion according to the manufacturer’s instructions.1 μg of total RNA from each sample was used for RNA-seq. A differential gene expression (DEG) [21] and differential alternative splicing (DAS) expression analysis was performed by GATC Biotech (Inview Transcriptome Discover, GATC Biotech AG, Constance, Germany) as previously described [31]. Briefly, for DAS analysis the aligned reads were used by multivariate analysis of transcript splicing (MATS) to detect alternative splicing events. MATS is a Bayesian statistical framework which uses a multivariate uniform prior to model the between-sample correlation in exon splicing patterns, and a Markov chain Monte Carlo method coupled with a simulation-based adaptive sampling procedure to calculate the p-values and false discovery rates (FDR) of differential alternative splicing [31]. A P_adjusted_<0.05 (FDR adjusted P-value) was used as a cut-off for differently alternative splicing events (DAS). DASs above the cutoff were analyzed for enrichment of gene ontology (GO) terms and pathways using database for annotation, visualization and integrated discovery (DAVID) v6.8 (Huang da, Sherman, & Lempicki, 2009a, 2009b). Default settings of DAVID were chosen except that the background database was restricted to the pool of genes annotated in our RNA-seq analysis [21]. Only gene enriched terms with a −log_10_ P-value<3 (p_value_<0.001) were considered significant. Delta “percent spliced in” (ΔPSI) distribution for two groups considered only DAS events detectable in both conditions tested.

### Data and statistical analysis

Each set of experiments contained mice injected with control or experimental viruses and were randomized per cage (i.e., each cage of four mice contained mice injected with control or experimental viruses). After stereotaxic surgery and until the end of each experiment, the experimenter was blind to the identity of the virus injected into each mouse. Behavioral experiments were performed three weeks after stereotaxic delivery of rAAVs. Gene enrichment analysis was performed using Fisher’s exact test P<0.001. Cumulative analysis was performed using paired two-tailed Student’s t test or Wilcoxon test, for normally and non-normal distributed data, respectively, p-values are shown on top of each panel. Statistics were performed using GraphPad prism for Mac OS X, version 8.

### Gene expression omnibus (GEO) accession

The RNA-seq data for alternative splicing analyzed in this study is publicly available at the National Center for Biotechnology Information (NCBI) Gene Expression Omnibus (GEO) with the accession number GSE107004.

## Results

### Spatial learning induces alternative splicing events that are altered in MeCP2 knockdown mice

In this study, we sought to investigate whether MeCP2 regulates alternative splicing events, in the adult hippocampus in basal conditions as well as after spatial learning. To this end, we delivered recombinant adeno-associated viruses (rAAV) into the adult dorsal hippocampus encoding either a control (Control-) or a MeCP2-specific shRNA sequence (Figure 1). We knocked down MeCP2 in neurons by using an AAV serotype (rAAV1/2) that displays predominant neuronal tropism [32, 33]. The viral construct also contained an expression cassette for mCherry under the control of the CamKIIα promoter (Figure 1) that served as an infection marker. This strategy allowed us to investigate MeCP2 function in the adult hippocampus without confounds originating from impaired postnatal neurodevelopment. We previously confirmed that this tool significantly decreases MeCP2 mRNA and protein levels selectively in the hippocampus. Moreover MeCP2-shRNA mice displayed impairments in hippocampus-dependent long term memory without exhibiting broad neurological impairments, such as motor deficits or anxiety-like behavior [21] that typically occur in animal models with disrupted MeCP2 expression from early developmental stages. Thus, this experimental strategy was chosen for gene expression analysis. In this experiment, half of the mice were kept in their homecage (baseline), whereas the remaining underwent spatial object location training (learning) (Figure 1). 30 min after the end of the task, a time point with robust transcriptomic changes after learning [21], we performed genome-wide differential alternative splicing analysis of the mouse dorsal hippocampal tissue expressing either MeCP2-shRNA or Control-shRNA in baseline conditions and after learning. RNA-seq analysis allowed the identification of five distinct mRNA splicing events: alternative 3′splice sites (A3SS), alternative 5′ splice sites (A5SS), mutually exclusive exons (ME),intron retention (IR) and exon skipping (ES) events (Figure 1).

**Figure 1.**
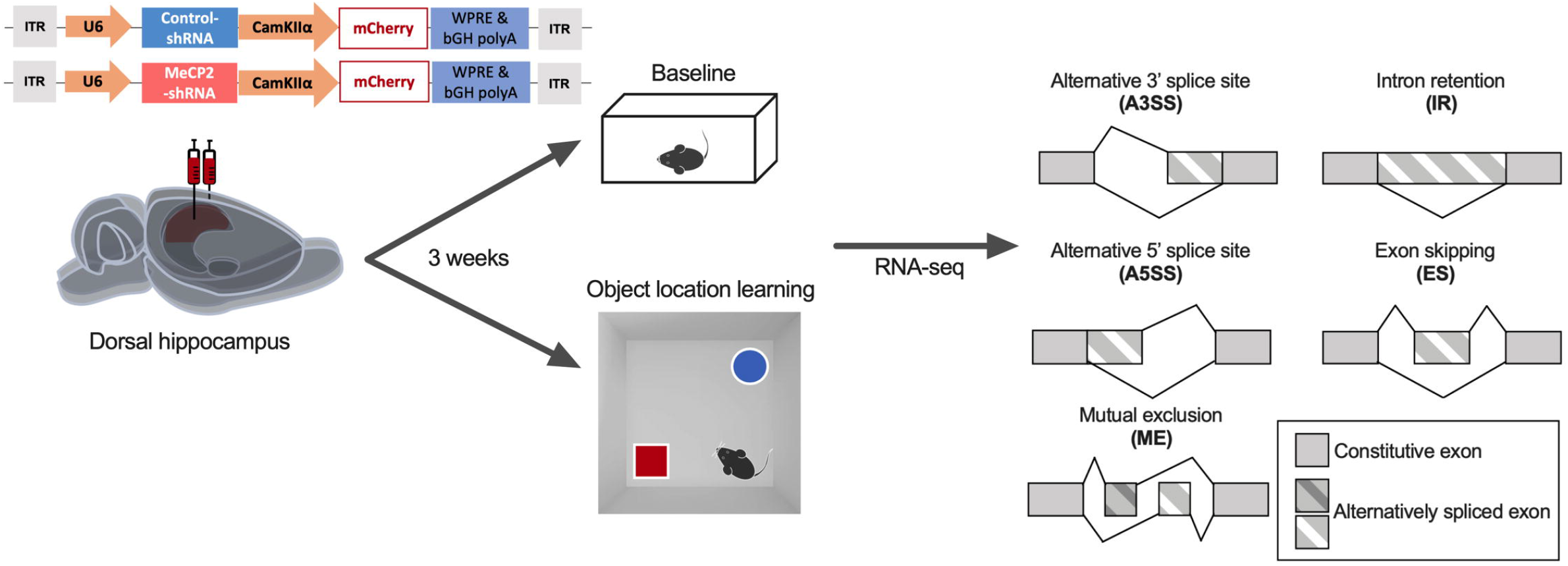
Schematic representation of the experimental design and alternative splicing events analyzed in this study. The viral vector contains a U6 promoter driving expression of MeCP2-shRNA or a control-shRNA and mCherry expression under the CamKIIα promoter. Three weeks after the delivery of recombinant AAVs into the dorsal hippocampus, mice remained either in home-cage (Baseline) or trained in object location learning. 30 minutes after training mice dorsal hippocampi were micro dissected and RNA-seq and alternative splicing analysis was performed. bGH polyA: Bovine growth hormone polyadenylation signal. ITR: inverted terminal repeat, WPRE: Woodchuck Hepatitis virus post-transcriptional regulatory element.

We started by identifying the DAS events induced by spatial learning in control mice, and then asked whether MeCP2 knockdown alters this learning-induced alternative splicing program. To determine this, we compared the alternative splicing profile of each treatment condition (Control- or MeCP2-shRNA) in basal conditions versus after learning (Figure 2A). We observed that object location learning induced 28 differential alternative splicing events in Control-shRNA-injected mice hippocampi, that consisted predominantly of ES events (42.8%) followed by A5SS (21.4%) and IR (17.9%), A3SS (17.9%) while no ME events were detected (Figure 2B) [see Additional file 1]. Some of the genes identified here have been previously described to undergo alternative splicing during memory consolidation or recall in a contextual fear conditioning paradigm (*Dnajb5* and *March7, Zfp207, Gls, Fuz*, respectively) [9]. In contrast, in MeCP2-shRNA expressing hippocampi, 13 learning-triggered DAS events were detected (Figure 2B).Furthermore, MeCP2-shRNA mice showed a clear shift towards more IR events (53.8%) and a reduced occurrence of ES (23.1%) and ME (7.7%), A5SS (7.7%) and A3SS (7.7%) in response to learning compared to the controls (Figure 2B) [see Additional file 1]. These findings indicate that MeCP2 reduction impaired DAS events in the adult hippocampus in response to spatial learning. Next, we analyzed whether there is a change in the fraction of the included or excluded isoforms in Control- or MeCP2-shRNA expressing mice using the delta “percent spliced in” (ΔPSI). The ΔPSI represents the difference between the ratio of transcripts that retain an intron/exon in relation to the total number of transcripts coded by a particular gene in two conditions. A ΔPSI value above or below 0% indicates an increased or reduced inclusion of alternative introns/exons, respectively. This parameter allows to investigate whether MeCP2 regulates the inclusion of introns/exons in alternatively spliced transcripts. Although MeCP2 reduction altered inclusion (ΔPSI>0) and exclusion (ΔPSI<0) events of each splicing subtype (Figure 2C and Supplementary Figure 1A-D), we focused on IR and ES events shown in Figure 2D since the majority of DAS belonged to these splicing categories, and MeCP2 induced an ES-IR switch. We found that MeCP2-shRNA animals showed a mild shift towards excluded IR events (14.3% included vs. 85.7% excluded) compared to controls (20% included vs. 80% excluded) and a decrease of ES (control: 16.7% included vs 83.3% excluded; MeCP2-shRNA: 33.3% included vs 66.7%. excluded) (Figure 2D) [see Additional file 2]. Similarly, hippocampal knockdown of MeCP2 led to alterations on A3SS, A5SS and ME inclusion/exclusion events (Supplementary Figure 1A-D). The majority of splicing events occurred in the same direction, that is inclusion or exclusion, in both control and MeCP2-shRNA animals (Figure 2E) [see Additional file 2]. Nonetheless, we also detected a subset of alternative splicing events that occurred in opposite directions, meaning that they underwent increased inclusion in MeCP2-shRNA mice and increased exclusion in Control-shRNA mice or vice versa (e.g. *Gls, Osmr, Trmt1, Irf7*) [see Additional file 2]. Remarkably, only 2 of the 13 DAS events observed in MeCP2-shRNA mice overlapped with the DAS events detected in Control-shRNA mice. This indicates that in MeCP2 knockdown conditions DAS events that occur in control conditions were no longer present (e.g. *Zmynd8, Nr3c1*) and new spliced isoforms were generated (11 unique DAS; e.g. *Atl2, Fhl1*) (Figure 2E). None of these events (neither overlapping nor unique) showed a bias towards any particular splicing type (Supplementary Figure 1E-I). Statistical analysis of all the DAS events detected in Control- or MeCP2-shRNA mice in response to learning did not show a statistically significant difference between the groups (Figure 2F and Supplementary Figure 1J-K).

**Figure 2.**
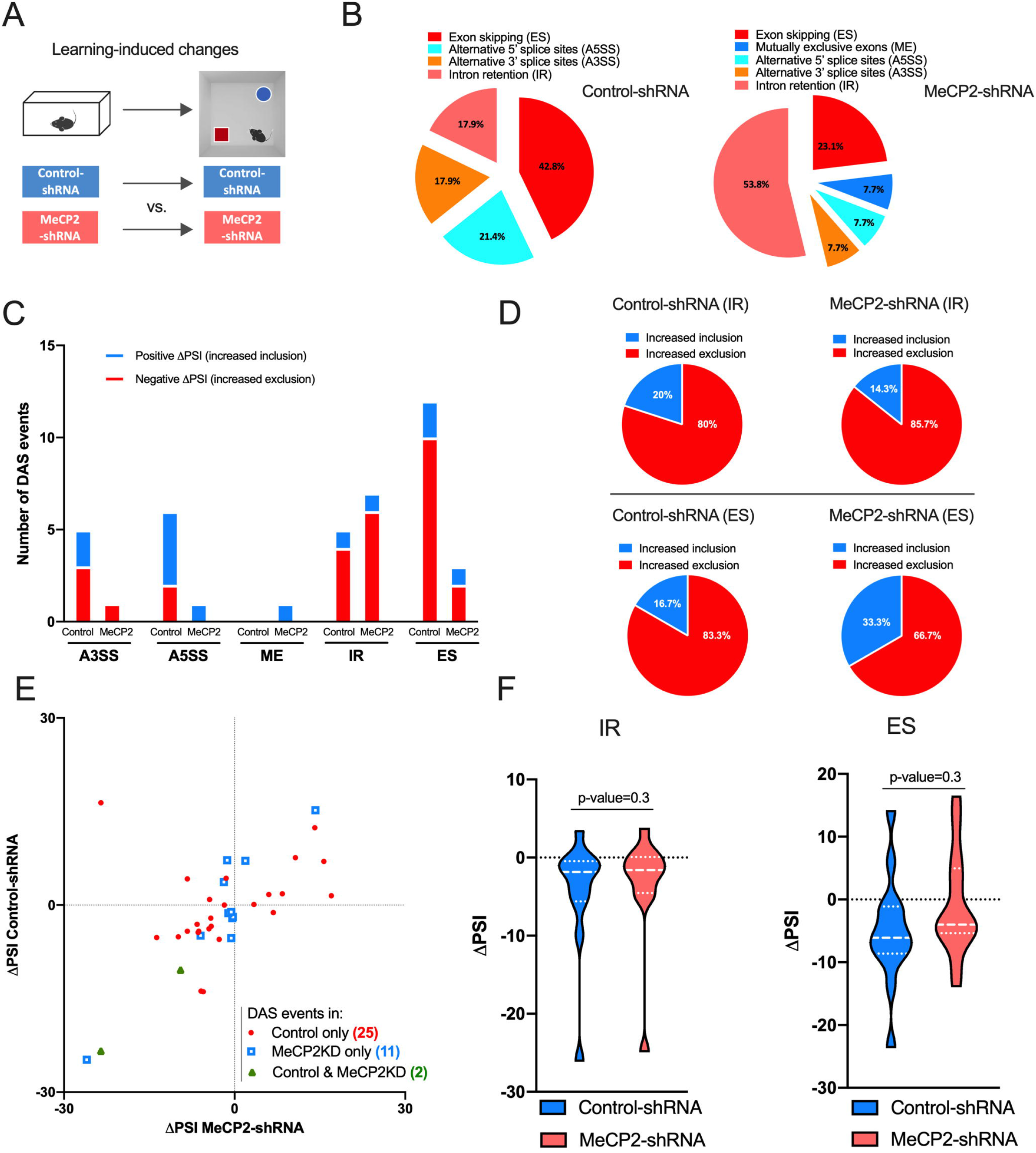
Spatial learning induces alternative splicing events that are altered in MeCP2 knockdown mice. A) Schematic representation of the comparisons used. B) Proportion of each differential alternative splicing events (DAS) in control conditions (left) or in MeCP2 knock-down conditions (right) in learning-induced conditions. C) Number of inclusion (positive ΔPSI, blue) and exclusion (negative ΔPSI, red) events for each type of alternative splicing modality in Control-shRNA (Control) and MeCP2-shRNA (MeCP2) mice (q-value<0.05). D) Pie charts showing the proportion of inclusion and exclusion events for intron retention (IR) and exon skipping (ES) in Control-shRNA and MeCP2-shRNA mice. E) Scatter plots showing changes in IR and ES events in Control-shRNA (Control) and MeCP2-shRNA (MeCP2KD) mice upon learning. Red dots and blue squares represent alternative splicing events that occurred in either Control or MeCP2 knockdown hippocampi (q-value<0.05), respectively. Green triangles represent alternative splicing events that occurred in both conditions (q-value<0.05). F) Violin plots showing the ΔPSI distribution of IR (left) and ES (right) events in Control-shRNA and MeCP2-shRNA hippocampi after learning. The P-values are based on paired two-tailed Student’s t test or Wilcoxon test and are indicated at the top of each panel. ΔPSI: delta “percent spliced in”.

To gain further insight into the functional categories of identified learning-induced DAS, we performed gene ontology (GO) analysis. For this, we carried two separate analysis for inclusion and exclusion DAS events (Figure 3A-B). We found that in control animals that underwent learning, GO terms associated with “Dendritic spine” and “Positive regulation of spine development” were predominantly enriched in the inclusion group (−log_10_P value<3), whereas terms associated with “Alternative splicing” and “Splice variant” showed a non-significant trend for enrichment in the exclusion cohort (−log_10_P value<3) [see Additional file 3]. These findings suggest that learning-induced alternative splicing events in the hippocampi of control mice are associated with dendritic spine regulation. Notably, in MeCP2-shRNA mice, there was no enrichment detected for the inclusion group, and the exclusion DAS cohort showed a non-significant trend for enrichment for terms associated with “Methylation”, “Splice variant”, “Alternative splicing” and “Compositionally bias region: Arg/Ser-rich” [see Additional file 3]. This data suggests that MeCP2 reduction compromises predominantly the learning-triggered processes associated with dendritic function.

**Figure 3.**
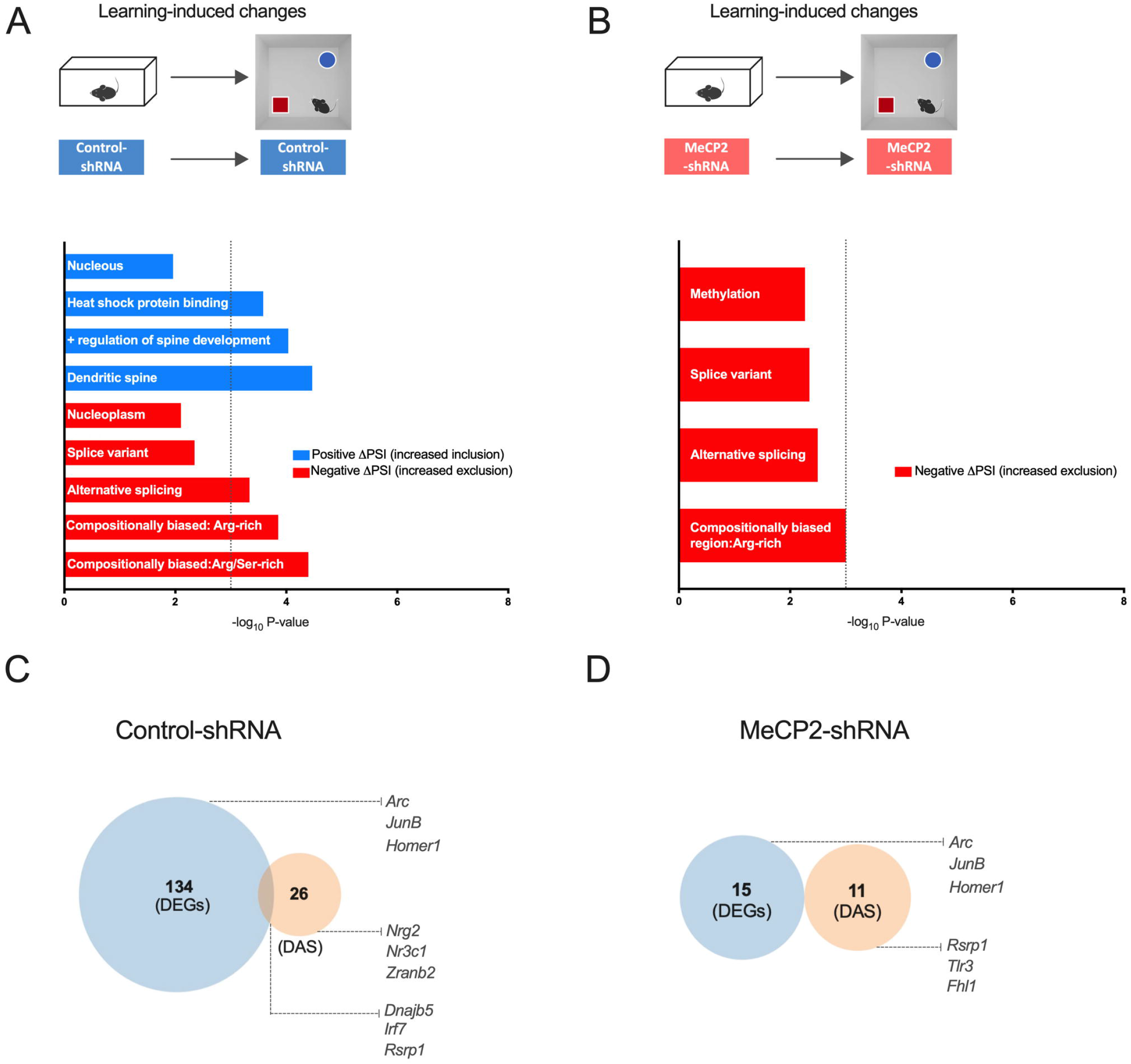
Analysis of genes that underwent differential alternative splicing events upon spatial learning. (A-B) Schematic representation of comparisons used (top). Gene ontology (GO) analysis for genes that underwent differential alternative splicing in the dorsal hippocampi of Control-shRNA (A) and MeCP2-shRNA (B) mice upon learning. Enriched GO terms (Fisher’s exact test P<0.001) for genes that underwent inclusion or exclusion (q-value < 0.05) events, upon learning. The blue and red bars represent −log_10_ (P-value) of the GO enrichment for inclusion and exclusion events, respectively. The vertical dashed line serves as a marker for P-value = 0.001 [−log_10_ (P-value) =3]. Absence of a colored bar means that genes of that GO term were not enriched in that specific category. C) Venn diagram showing overlap between total number of differentially expressed genes (DEGs) and genes that underwent differential alternative splicing events (DAS) in home-cage (baseline) conditions when MeCP2 was knocked down in the adult dorsal hippocampus. D) Venn diagram showing overlap between total number of differentially expressed genes and genes that underwent differential alternative splicing events in learning state (learning) conditions when MeCP2 was knocked down in the adult dorsal hippocampus.

Considering that MeCP2 is required for optimal expression and alternative splicing of several genes, we asked to which degree these two gene populations (DEGs and DASs) overlap. This analysis indicates whether MeCP2 uses these two regulatory mechanisms on similar or different genes. Since the same tissue was used for DAS and for the previously published differential gene expression analysis [21], the two datasets could be directly compared. To this end, we identified genes that underwent alternative splicing, and compared this gene population to the differentially expressed genes (DEGs) in the same conditions (learning-induced DEGs were compared to learning-induced DASs in Control- or MeCP2-shRNA expressing mouse hippocampus) (Figure 3C,D). We found that in control group, only 3 DEGs showed also DAS events in response to learning (out of 134 DEGs and 26 DAS) (Figure 3C and Additional file 3). In MeCP2-shRNA animals, the differentially expressed genes in response to learning and the learning-induced differential alternative splicing events did not overlap (Figure 3D and Additional file 3). Taken together, this data indicates that learning induces changes in the expression levels and in the predominance of specific alternatively spliced variants of distinct gene populations. Furthermore, our results implicate a requirement for MeCP2 in both processes.

### MeCP2 knock-down changes splicing events in baseline and learning states

To gain a deeper understanding of how MeCP2 regulates alternative splicing events, we asked whether acute MeCP2 reduction influences DAS events already in baseline and/or after learning conditions. To this end, we compared the alternative splicing profile of Control-versus MeCP2-shRNA mice in basal conditions, as well as after learning (hereafter, learning state) (Figure 4A). We identified a total of 156 DAS events (q-value < 0.05) in baseline conditions upon MeCP2 disruption in the hippocampus [see Additional file 4]; ES events were predominant (75%), followed by IR (10.3%), ME (6.4%), A5SS (4.5%) and A3SS (3.8%) (Figure 4B). Altered alternative splicing in overlapping genes has been observed in the hippocampus of *Mecp2-null* mice (i.e. *Zfp207, Map4* and *Ppfia2*) [8]. Similarly, DAS profile of MeCP2-shRNA hippocampus after learning was different from the controls. We identified 94 DAS events (q-value < 0.05) in MeCP2-shRNA mice in learning state [see Additional file 4], in which ES events were predominant (70.2%), followed by IR (25.5%) and A3SS (4.3%) whereas no A5SS and ME events were detected (Figure 4B). Next, we determined the change in the fraction of the included or excluded events of each splicing subtype in baseline or learning in MeCP2-knockdown conditions (Figure 4C-D). We found that MeCP2-shRNA animals have preferentially decreased IR in baseline conditions (31.2% inclusion vs. 68.8% exclusion) while in learning state, the relative abundance of inclusions/exclusions in MeCP2-shRNA mice shifted predominantly towards included introns (66.7% inclusion vs. 33.3% exclusion) (Figure 4D) [see Additional file 5]. Interestingly, although the total number of ES events in MeCP2-disrupted hippocampus decreased by learning (Figure 4C), the proportion of inclusions/exclusions among the total ES events remained similar in baseline and learning state (baseline: 32.5% included vs. 67.5% excluded; learning 37.9%: vs. 62.1%) (Figure 4D). Similarly, hippocampal knockdown of MeCP2 lead to alterations on A3SS, A5SS and ME inclusion/exclusion events in baseline and in learning conditions (Supplementary Figure 2A-D) [see Additional file 5]. Next, we checked the common DAS events in baseline and learning state in MeCP2-shRNA conditions. We found that, in baseline and in learning state, hippocampal MeCP2 reduction led to 131 and 75 unique DAS events, respectively (Figure 4E). Only 19 DAS events occurred in both conditions, suggesting that learning induces distinct alternative splicing events. Notably, the majority of DAS events detected in baseline or after learning happened in the same direction in MeCP2-disrupted and control hippocampi, although a small subset of splicing events occurred in opposite ways (Figure 3E and Additional file 5). Deeper analysis revealed that the oppositely regulated DAS subset showed no bias for a particular splicing event type (Supplementary Figure 2E-I) [see Additional file 5]. Cumulative analysis of all DAS events in MeCP2-disrupted hippocampus showed a significant increase in retained introns (Figure 3F), a decrease in skipped exons (Figure 4F) and either a significant or a non-significant trend for increase of A5SS and ME inclusion events in learning state compared to baseline (Figure 4F and Supplementary Figure 2J-L). No change was detected for A3SS events (Supplementary Figure 2J). These results show that MeCP2 reduction induces a distinct DAS profile in baseline and upon learning. Thus, indicating that the differences found in the learning state do not only reflect changes in basal conditions, but also a requirement for MeCP2 in learning-dependent alternative splicing.

**Figure 4.**
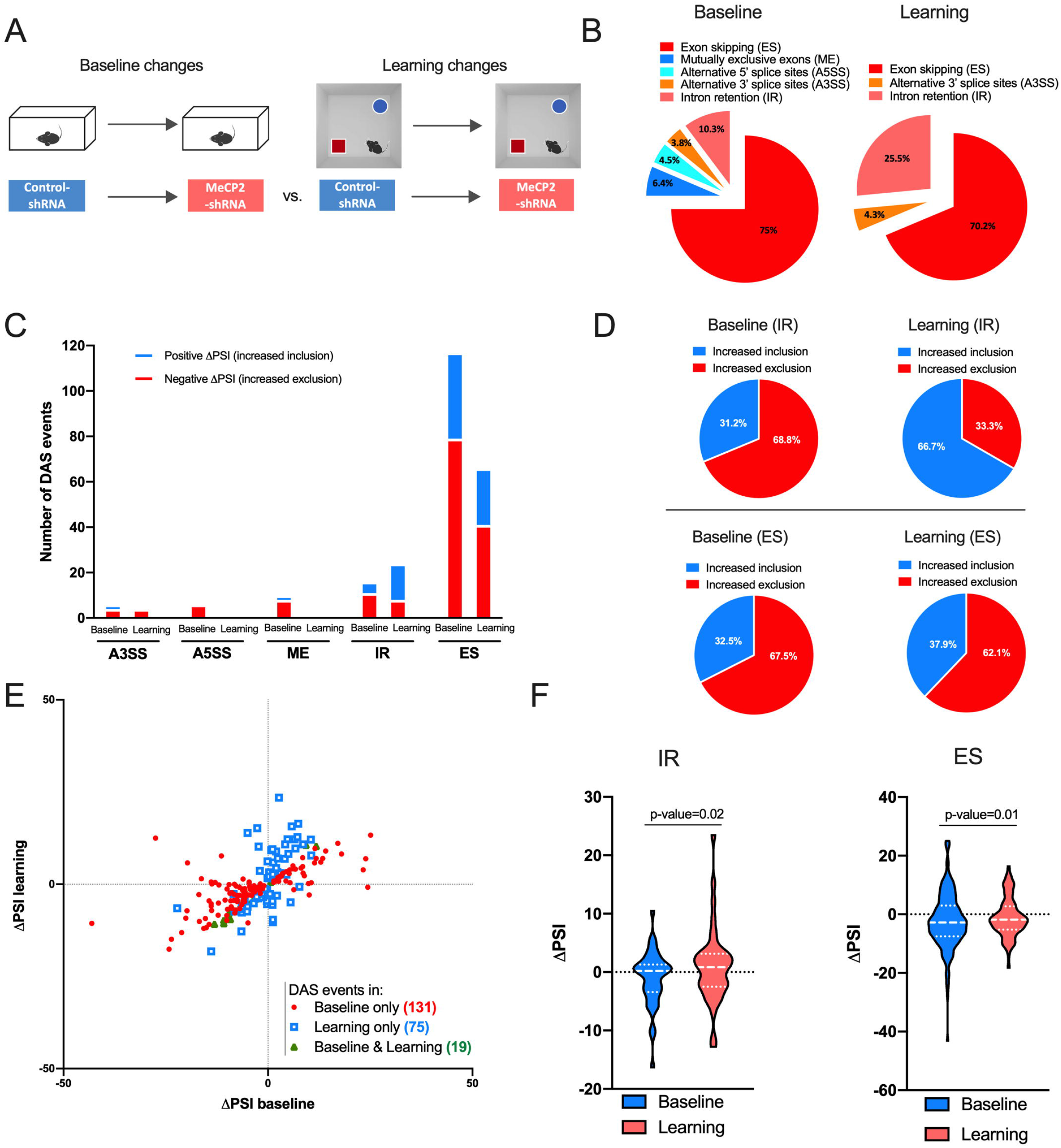
MeCP2 knockdown changes baseline and learning-associated splicing events. A) Schematic representation of the comparisons used. B) Proportion of each differential alternative splicing events (DAS) in baseline conditions (left) or in learning state (right) in MeCP2-shRNA conditions. C) Number of inclusion (positive ΔPSI, blue) and exclusion (negative ΔPSI, red) events for each type of alternative splicing modality in MeCP2 knockdown during baseline and learning state conditions (q-value<0.05). D) Pie charts showing the proportion of inclusion and exclusion events for intron retention (IR) and exon skipping (ES) in baseline and learning state conditions. E) Scatter plots showing changes in IR and ES events in home-cage (Baseline) and learning state (learning) conditions by MeCP2 knockdown. Red dots and blue squares represent alternative splicing events occurred in either baseline or learning state (q-value<0.05) conditions, respectively. Green triangles represent alternative splicing events that occurred in both conditions (q-value<0.05). F) Violin plots showing the ΔPSI distribution of IR (left) and ES (right) events in baseline and learning state in the hippocampi of MeCP2-shRNA mice. The P-values are based on paired two-tailed Student’s t test or Wilcoxon test and are indicated at the top of each panel.

Next, to gain insight into the functional categories of the genes that required MeCP2 for alternative splicing in baseline or learning states, we performed GO analysis. This was applied to both conditions (baseline or learning) and were divided into inclusion (ΔPSI>0) and exclusion (ΔPSI<0) events. We found that DAS inclusions in MeCP2 reduction in baseline conditions were enriched for terms such as “Phosphoprotein”, “Alternative splicing” and “Cytoskeleton”, whereas DAS exclusions in MeCP2-shRNA mice were associated with the functional categories termed “Alternative splicing”, “Clathrin vesicle coat”, “Tubulin binding” (−log_10_P value<3) (Figure 5A) [see Additional file 6]. After learning, only enrichment for “Alternative Splicing” for inclusion events and “Cell-cell adherent junctions”, “Neuronal cellular homeostasis” and “Positive regulation of protein binding” for increased exclusion events were found (Figure 5B) [see Additional file 6]. These results indicate that both in baseline and after learning conditions MeCP2 regulates DAS events associated with general neuronal function processes despite that DAS events are generally distinct in both conditions (Figure 4E-F).

**Figure 5.**
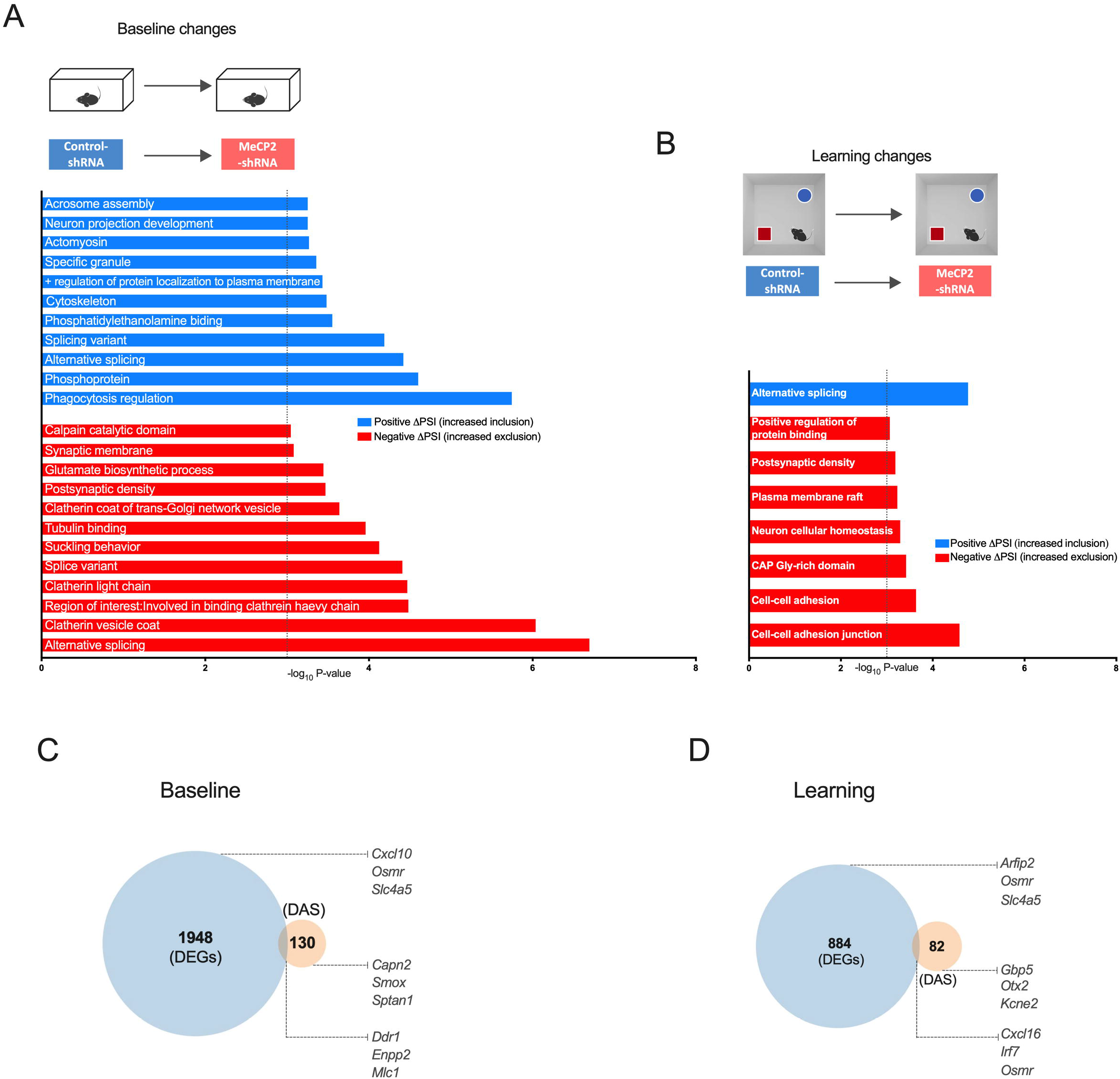
Analysis of genes that underwent differential alternative splicing events in baseline and in learning state upon MeCP2 knock-down. (A-B) Schematic representation of comparisons used (top). Gene ontology (GO) analysis for genes that underwent differential alternative splicing in the dorsal hippocampi MeCP2-shRNA mice in baseline (A) and learning state (B) conditions. Enriched GO terms (Fisher’s exact test P<0.001) for genes that underwent inclusion or exclusion (q-value < 0.05) events. The blue and red bars represent −log_10_ (P-value) of the GO enrichment for inclusion and exclusion events, respectively. The vertical dashed line serves as a marker for P-value =0.001 [−log_10_ (P-value) =3]. Absence of a colored bar means that genes of that GO term were not enriched in that specific category. ΔPSI: delta “percent spliced in”. C) Venn diagram showing overlap between total number of differentially expressed genes and genes that underwent differential alternative splicing events in learning-induced conditions in the adult dorsal hippocampus of control mice (control-shRNA). D) Venn diagram showing overlap between total number of differentially expressed genes and genes that underwent differential alternative splicing events in learning-induced conditions when MeCP2 was knocked down in the adult dorsal hippocampus (MeCP2-shRNA).

Next, we compared DEGs and DASs in MeCP2-reduced hippocampi in baseline or learning states. We found that only 17 differentially expressed genes in MeCP2 knockdown also showed altered alternative splicing (out of 1948 DEGs and 130 DAS) in baseline conditions (Figure 5C and Additional file 6), whereas this number was as low as 7 genes in learning state (out of 884 DEGs and 82 DAS) (Figure 4B and Additional file 6). Altogether, these findings indicate that MeCP2 regulates the predominance of specific alternatively spliced variants mostly without affecting the overall level of transcripts coded by that gene both in baseline conditions and after learning.

## Discussion

In this study, we showed that adult hippocampal MeCP2 is required for the regulation of alternative splicing events during memory consolidation. We demonstrated that MeCP2 preserves the alternative splicing profile of mature hippocampal neurons and regulates learning-dependent splicing of genes important for neuronal structure and function. Therefore, our findings show that MeCP2 not only regulates the levels of expression of memory-related genes, but also the relative abundance of specific alternatively spliced isoforms, thus uncovering another mechanism by which MeCP2 impacts neuronal functional and structural properties during memory consolidation. This highlights a multifactorial requirement for MeCP2 in adult cognitive processes.

MeCP2 has well-established functions during neurodevelopment as evidenced by the severe neurological impairments characteristic of RTT, a neurodevelopmental disorder caused by mutations in the *Mecp2* gene [18, 20]. Furthermore, several lines of evidence also support an important function during adulthood; MeCP2 is expressed at high levels in the adult brain [34] and is required for its function [21, 34–39]. Specifically, it has been demonstrated that MeCP2 plays an important role in adult cognitive abilities [18, 21]. Mounting evidence indicates that long-lasting synaptic remodeling important for memory consolidation is supported not only by learning-triggered changes in the transcriptional, but also in the post-transcriptional profile [9] of neurons. In this study, we investigated the regulatory function of MeCP2 in alternative splicing mechanisms. To this end, we selectively decreased MeCP2 levels in adult hippocampal neurons [21]. This way, we could dissect the impact of MeCP2 disruption on the alternative splicing profile of mature hippocampal neurons without confounds originating from impaired neurodevelopment. We found that reducing MeCP2 expression of mature hippocampal neurons led to aberrant alternative splicing profiles. This finding is in line with previous studies that demonstrated a role for MeCP2 in alternative splicing regulation [24, 26, 27, 40]. Several studies analyzed genome-wide gene expression changes in response to learning and have shown the requirement for MeCP2 for this learning-dependent gene expression [21, 41]. In contrast, alternative splicing changes on a genome-wide scale upon learning have been less explored. Poplawski and colleagues (2016) were the first to investigate genome-wide alternative splicing changes in the hippocampus after a contextual-fear learning and after memory recall and identified novel alternative splicing isoforms that may be critical for memory consolidation [9]. Our observations support and further expand this previous dataset by providing a novel set of alternative splicing events triggered by a non-aversive object-location learning, concluding that both aversive and non-aversive forms of learning induce genome-wide alternative splicing changes in the hippocampus.

The mechanisms through which MeCP2 regulate learning-dependent alternative splicing events, particularly in mature neurons, are poorly understood. Recently, Osenberg and colleagues (2018) studied activity-dependent gene expression and alternative splicing in a mouse model of RTT. The authors elicited neuronal activity in *Mecp2-null* (Mecp2^−/y^) mice through the administration of kainic acid and identified genome-wide alternative splicing changes in the hippocampus in response to this neuronal stimulation. They found an aberrant global pattern of gene expression and alternative splicing events. Here, we used an adult-onset knockdown of MeCP2 and induced neuronal activity by a physiological and memory-relevant stimulus, novel environment exposure. We found that MeCP2 knockdown led to an increase in intron retention and decreased excluded exons. Notably, Wong and colleagues (2017) showed that decreased MeCP2 binding near splice junctions facilitates intron retention via reduced recruitment of splicing factors, such as the splicing factor transformer-2 protein homolog beta (Tra2b), and stalling of RNA polymerase II [26]. In MeCP2 depletion conditions, like the one in our study, intron retention is favored possibly through the enabling of Tra2b activity. Importantly, this was not associated with an altered Tra2b expression in MeCP2-shRNA mice [21]. Moreover, intragenic DNA methylation and MeCP2 binding promote exon recognition and consequently MeCP2 ablation results in aberrant exon skipping events [25]. Overall, the demonstrated involvement of MeCP2 in these splicing modalities together with the shift towards increased retained introns and exons in MeCP2 knockdown conditions observed in our study, suggest that MeCP2 contributes to learning-induced alternative splicing through these mechanisms. Although aberrations in these splicing events were predominant, we identified learning-induced changes in other forms of alternative splicing in the hippocampus of MeCP2-shRNA mice. This indicates that MeCP2 may regulate other forms of splicing through mechanisms not yet identified.

In this study, we analyzed alternative splicing events in response to learning in control or MeCP2-shRNA hippocampi as well as in baseline or learning states. This combinatorial analysis allowed us to conclude that the differences found in the learning state do not only reflect changes in basal conditions, but also a requirement for MeCP2 in learning-dependent alternative splicing. Therefore, this indicates that the contribution of MeCP2 to synaptic plasticity and memory is likely two-fold. On the one hand, MeCP2 regulates the neuronal basal transcriptome which may impact neuronal properties such as synaptic transmission and intracellular signal transduction, and additionally may regulate directly stimulus-dependent transcriptional and post-transcriptional events in the nucleus.

MeCP2 is essential for the maintenance of structural integrity of neuronal circuits as demonstrated in RTT mouse models that show defects in dendritic structure and arborization [42–44]. We found that upon learning, genes associated with “Positive regulation of spine development” and “Dendritic spine” showed an enrichment in control conditions, but not in MeCP2 disruption. Noteworthy examples are the nuclear receptor subfamily 3, group C, member 1 (*Nr3c1*) and the zinc finger, MYND-type containing 8 (*Zmynd8*) that play important roles in the regulation of cellular responses to stress and the formation of dendritic spines, respectively. *Nr3c1* encodes for the glucocorticoid receptor (GR) which upon binding of glucocorticoids affects a wide range of processes including gene expression, synaptic plasticity, dendritic morphology and learning and memory [45, 46]. The gene is composed of 9 exons and several isoforms are generated through alternative splicing [47, 48]. These isoforms have opposite binding affinities for glucocorticoids, distinct cellular localization and gene activation properties [47]. A tight control of the amounts of different alternatively spliced GR isoforms is therefore crucial for proper cellular function. Our data showed that learning-induced increase in an A3SS isoform of this receptor is disrupted in MeCP2-knockdown animals. Splicing in this region is thought to regulate the expression of different isoforms which may impact cognition [48]. The *Zmynd8* (or *Spikar*) gene is expressed at high levels in the brain and the expression of its splicing variants depend on neuronal activity [49]. Spikar is present at dendritic spines and when knocked down reduces the density of dendritic spines in cultured hippocampal neurons leading to decreased excitatory synapses [49]. In our study, control animals showed an increase in a particular spliced version (A5SS) of *Spikar* that was no longer induced in the hippocampus of MeCP2-shRNA mice.

Altogether this data suggests that alterations in the relative amounts of splicing isoforms of genes supporting structural plasticity changes after learning may contribute to the cognitive deficits observed in these mice [21]. It is noteworthy that acute disruptions of adult hippocampal MeCP2 did not alter the dendritic complexity and spine density of CA1 neurons in baseline conditions [21]. This is in line with our observations that DAS in MeCP2 knockdown in baseline conditions was not enriched for genes functionally relevant to “dendritic spine regulation”. Our findings therefore suggest that MeCP2 regulates alternative splicing of the genes associated with dendritic spines mostly in response to learning, which may cause selective impairments in learning-dependent spine remodeling [50–52]. Whether MeCP2 disruptions alter learning-dependent structural remodeling in mature hippocampal neurons remains to be investigated.

We found that at baseline conditions MeCP2 reduction promoted an overall increase in IR and a decrease in skipped exons, particularly in genes functionally linked to general neuronal functions. Specifically, the abundance of spliced isoforms relevant for neurotransmitter synthesis (glutaminase (*Gls*)), vesicle recycling (synaptojanin 1 (*Synj1*)) and neurotransmitter receptors (gamma-aminobutyric acid (GABA) A receptor, subunit gamma 2 (*Gabrg2*), glutamate ionotropic receptor NMDA Type Subunit 1 (*Grin1*)) was altered in MeCp2 knockdown conditions. Interestingly, the *Grin1* gene gives rise to 8 splice variants and recently it has been shown that the selective expression of different GluN1 isoforms determines long-term potentiation in the hippocampus and spatial memory performance [53]. Moreover, the relative abundance of some spliced isoforms of GluN1 subunit is associated with increased seizure susceptibility in adult mice [54]. Taken together, these findings suggest that altered alternative splicing events observed in MeCP2-shRNA mice at baseline might impact proper neuronal function and consequently contribute to cognitive deficits and excitation/inhibition imbalance reminiscent of RTT. Furthermore, we found aberrant splicing and/or expression of splicing regulators in resting and learning conditions. In particular, MeCP2-shRNA mice during the learning state displayed changes in the abundance of U1 small nuclear ribonucleoprotein 70 (Snrnp70) and U2 small nuclear RNA auxiliary factor 1-like 4 (U2af1l4) spliced variants, two components of the spliceosome. In baseline conditions, MeCP2 regulates the expression of the Small the Nuclear Ribonucleoprotein U4/U6.U5 Subunit 27 (Snrnp27) and the Polypyrimidine tract-binding protein 1 (Ptbp1) [21]. These findings are in agreement with a previous study that also observed alterations in the expression and splicing of splicing regulators as a consequence of MeCP2 ablation [8]. It is plausible that aberrant expression and/or splicing levels of splicing mediators may induce a second wave of impairments in downstream splicing events, such as in response to learning as observed in MeCP2-shRNA mice. Furthermore, as MeCP2 interacts not only with transcription factors but also with regulators of alternative splicing [24–27, 40, 55], loss of MeCP2 may thus impair their recruitment and promote the disruption of alternative splicing events observed in MeCP2-shRNA mice.

Overall in this study, we found that spatial learning induces alternative splicing events of transcripts with relevant functions for neuronal structure and function. Moreover, our findings implicated MeCP2 in the regulation of this process. We showed that the reduction of MeCP2 levels in adult hippocampus promoted aberrant alternative splicing patterns both in baseline and learning states. This study uncovered another factor that likely contributes to the neuronal dysfunctions that characterize RTT.

## List of abbreviations

A3SS: Alternative 3′ splice sites
A5SS: Alternative 5′ splice sites
DAS: Alternative splicing events
DAVID: Database for annotation, visualization and integrated discovery
DEG: Differentially expressed gene
ES: Exon skipping
FDR: False discovery rate
Gabrg2: gamma-aminobutyric acid (GABA) A receptor, subunit gamma 2
GEO: Gene Expression Omnibus
Gls: Glutaminase
GO: Gene ontology
GR: Glucocorticoid receptor
Grin1: Glutamate Ionotropic Receptor NMDA Type Subunit 1
IR: Intron retention
Luc7l3: LUC7 Like 3 Pre-MRNA Splicing Factor
ME: Mutually exclusive exons
MeCP2: Methyl CpG binding domain protein 2
NCBI: National Center for Biotechnology Information
Nr3c1: Nuclear receptor subfamily 3, group C, member 1
rAAV: Recombinant adeno-associated virus
RTT: Rett Syndrome
shRNA: short-hairpin RNA
Snrnp70: U1 small nuclear ribonucleoprotein 70
Synj1: synaptojanin 1
Tra2b: Transformer-2 protein homolog beta
U2af1l4: U2 small nuclear RNA auxiliary factor 1-like 4
YB-1: Y-box binding protein 1
Zmynd8: Zinc finger, MYND-type containing 8
ΔPSI: percent spliced in

## Declarations

### Ethics approval

All animal experiments were done in accordance with German guidelines for the care and use of laboratory animals and with the European Community Council Directive 2010/63/EU. Experiments were approved by local authorities (Regierungspraesidium Karlsruhe, Germany).

### Consent for publication

Not applicable.

### Availability of data and materials

The datasets generated and analyzed during the current study are available at the National Center for Biotechnology Information (NCBI) Gene Expression Omnibus (GEO) with the accession number GSE107004 and are included in the additional files.

### Authors contributions

D.V.C.B., K.G.K. and A.M.M.O. conceived the project and designed research; D.V.C.B. and K.G.K. performed research; D.V.C.B. analyzed data; D.V.C.B., K.G.K. and A.M.M.O. wrote the manuscript.

## Acknowledgements

We thank to Benjamin Zeuch for producing the viruses and providing technical assistance throughout the project and to Janina Kupke and Lukas Frank for comments to the manuscript.

---

**Supplementary Figure 1.**
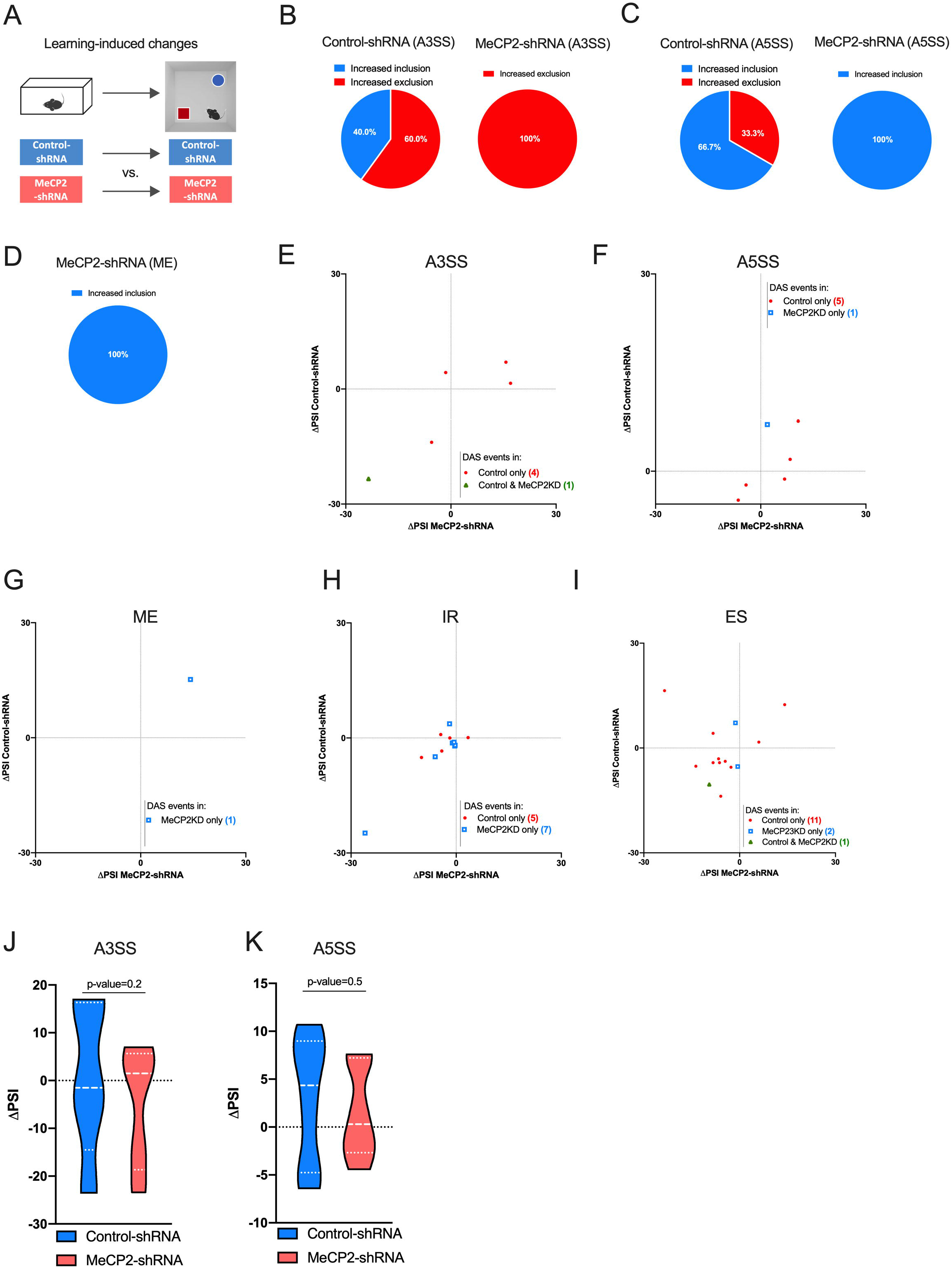
Alternative splicing event-specific changes in MeCP2 knockdown mice upon spatial learning. A) Schematic representation of the comparisons used. B-D) Pie charts showing the proportion of learning-induced inclusion and exclusion events for (B) alternative 3′ splice sites (A3SS), (C) alternative 5′ splice (A5SS) and (D) mutually exclusive exons (ME) in Control-shRNA and MeCP2-shRNA mice. E-I) Scatter plots showing changes in (E) A3SS, (F) A5SS, (G) ME, (H) intron retention (IR) and (I) exon skipping (ES) events in Control-shRNA (Control) and MeCP2-shRNA (MeCP2KD) mice upon learning. Red dots and blue squares represent alternative splicing events occurred in either Control or MeCP2-knock-down (MeCP2KD) (q-value<0.05), respectively. Green triangles represent alternative splicing events that occurred in both conditions (q-value<0.05). J-L) Violin plots showing the ΔPSI distribution of A3SS (J), A5SS (K) events in Control-shRNA and MeCP2-shRNA hippocampi after learning. The P-values are based on paired two-tailed Student’s t test or Wilcoxon test. test and are indicated at the top of each panel. ΔPSI: delta “percent spliced in”.

**Supplementary Figure 2.**
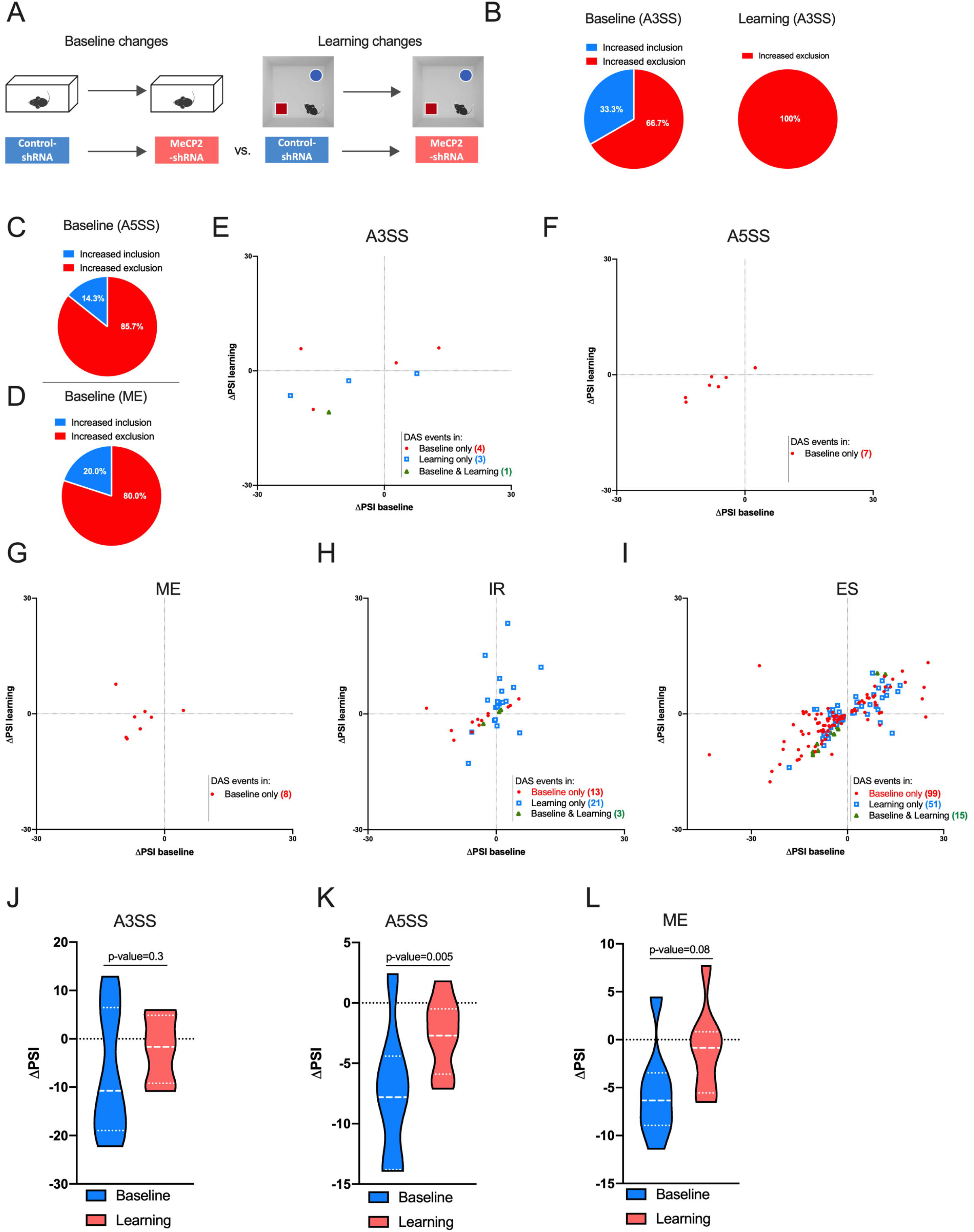
Alternative splicing event-specific changes in MeCP2 knock-down mice during baseline and learning state conditions. A) Schematic representation of the comparisons used. B-D) Pie charts showing the proportion of inclusion and exclusion events for (B) alternative 3′ splice sites (A3SS), and (C) alternative 5′ splice (A5SS) and mutually exclusive exons (ME) in MeCP2-shRNA mice during baseline and learning-state conditions. Note that no ME changes were detected by MeCP2 knockdown in learning-state. E-I) Scatter plots showing changes in (E) A3SS, (F) A5SS, (G) ME, (H) intron retention (IR) and (I) exon skipping (ES) events in home-cage (Baseline) and learning state (Learning) conditions in MeCP2-shRNA mice compared to the controls. Red dots and blue squares represent alternative splicing events occurred in either baseline or learning conditions (q-value<0.05), respectively. Green triangles represent alternative splicing events that occurred in both conditions (q-value<0.05). J-L) Violin plots showing the ΔPSI distribution of (J) A3SS, (K) A5SS and (L) ME events in baseline and learning state conditions in the dorsal hippocampi of MeCP2-shRNA mice. The P-values are based on paired two-tailed Student’s t test or Wilcoxon test. and are indicated at the top of each panel. ΔPSI: delta “percent spliced in”.

## Additional Files

- Additional file 1
  - File format: Microsoft Excel (.xlsx)
  - Title of data: Learning-induced DAS
  - Description of data: This file contains the number of differential alternative splicing (DAS) events per subtype for learning-induced conditions.
- Additional file 2
  - File format: Microsoft Excel (.xlsx)
  - Title of data: ΔPSI for Learning-induced DAS
  - Description of data: This file contains ΔPSI values for the differential alternative splicing (DAS) events per subtype for learning-induced conditions.
- Additional file 3
  - File format: Microsoft Excel (.xlsx)
  - Title of data: GO analysis and DEG/DAS overlap for Learning-induced conditions
  - Description of data: This file contains gene ontology (GO) analysis and overlap for DEGs and DAS for learning-induced conditions.
- Additional file 4
  - File format: Microsoft Excel (.xlsx)
  - Title of data: Baseline and learning state DAS
  - Description of data: This file contains the number of differential alternative splicing (DAS) events per subtype for baseline and learning state conditions.
- Additional file 5
  - File format: Microsoft Excel (.xlsx)
  - Title of data: ΔPSI for Baseline and learning state DAS
  - Description of data: This file contains ΔPSI values from the differential alternative splicing (DAS) events per subtype for baseline and learning-state conditions.
- Additional file 6
  - File format: Microsoft Excel (.xlsx)
  - Title of data: GO analysis and DEG/DAS overlap for baseline and learning state conditions
  - Description of data: This file contains gene ontology (GO) analysis and overlap for DEGs and DAS for baseline and learning conditions.

